# Species-specific adaptations determine how aridity and biotic interactions drive the assembly of dryland plant communities

**DOI:** 10.1101/147181

**Authors:** Miguel Berdugo, Fernando T. Maestre, Sonia Kéfi, Nicolas Gross, Yoann Le Bagousse-Pinguet, Santiago Soliveres

## Abstract

1. Despite being a core ecological question, disentangling individual and interacting effects of plant-plant interactions, abiotic factors and species-specific adaptations as drivers of community assembly is challenging. Studies addressing this issue are growing rapidly, but they generally lack empirical data regarding species interactions and local abundances, or cover a narrow range of environmental conditions.
2. We analysed species distribution models and local spatial patterns to isolate the relative importance of key abiotic (aridity) and biotic (facilitation and competition) drivers of plant community assembly in drylands worldwide. We examined the relative importance of these drivers along aridity gradients and used information derived from the niches of species to understand the role that species-specific adaptations to aridity play in modulating the importance of community assembly drivers.
3. Facilitation, together with aridity, was the major driver of plant community assembly in global drylands. Due to community specialization, the importance of facilitation as an assembly driver decreased with aridity, and became non significant at the border between arid and semiarid climates. Under the most arid conditions, competition affected species abundances in communities dominated by specialist species. Due to community specialization, the importance of aridity in shaping dryland plant communities peaked at moderate aridity levels.
4. Synthesis: We showed that competition is an important driver of community assembly even under harsh environments, and that the effect of facilitation collapses as driver of species relative abundances under high aridity because of the specialization of the species pool to extremely dry conditions. Our findings pave the way to develop more robust species distribution models aiming to predict the consequences of ongoing climate change on community assembly in drylands, the largest biome on Earth.

## Introduction

Climate change is affecting biodiversity by reducing local species richness, altering composition and homogenizing biotas in terrestrial ecosystems worldwide (Millenium Ecosystem Assessment 2005). A major challenge is to accurately predict future community composition and local species abundance to better understand how these changes will impact ecosystem structure and functioning (Chapin III et al. 2000; Valencia et al. 2015). However, predicting the local abundances of species is a difficult task because of the interplay between abiotic and biotic factors that joinly determine community composition and structure (Lawton 1999; Soberón 2007; Mayfield & Levine 2010). In adition, the relative importance of both biotic interactions and abiotic factors is highly dependent on species features such as their ability to cope with abiotic stress, making the outcomes of biotic interaction highly species-specific and context dependent (Choler, Michalet & Callaway 2001; Liancourt, Callaway & Michalet 2005; Gross et al. 2010; Boulangeat, Gravel & Thuiller 2012; Soliveres et al. 2014).

Combining approaches focusing on contrasting spatial scales, such as species distribution models (SDMs, applied at regional scales, Guisan & Thuiller 2005) and local spatial segregation and aggregation data (applied at local scales) can help to tease apart the relative importance of abiotic and biotic assembly drivers (Fig. 1). SDMs rely on abiotic variables, such as climate and soil type, to assess the probability of occurrences of species in a given site. They are able to predict diversity changes and extinction risks at regional scales, and account for the effect of abiotic factors (e.g., Araújo et al. 2002). Therefore, SDMs can help to identify the upper limit of species abundance within a community based on its environmental suitability (VanDerWal etal. 2009). Along with their ability to describe environmental suitability of individual species, SDMs allow to assess species adaptive strategies such as the degree of specialization to environmental conditions (Devictor et al. 2010), an important parameter to predict the outcomes of biotic interactions (Liancourt et al. 2005; Gross et al. 2010). Spatial co-occurrence patterns observed within communities, in turn, can inform about the frequency and strength of plant-plant interactions (e.g., Cavieres et al. 2006), but provide limited information about the role of abiotic factors on community assembly.

**Figure 1.**
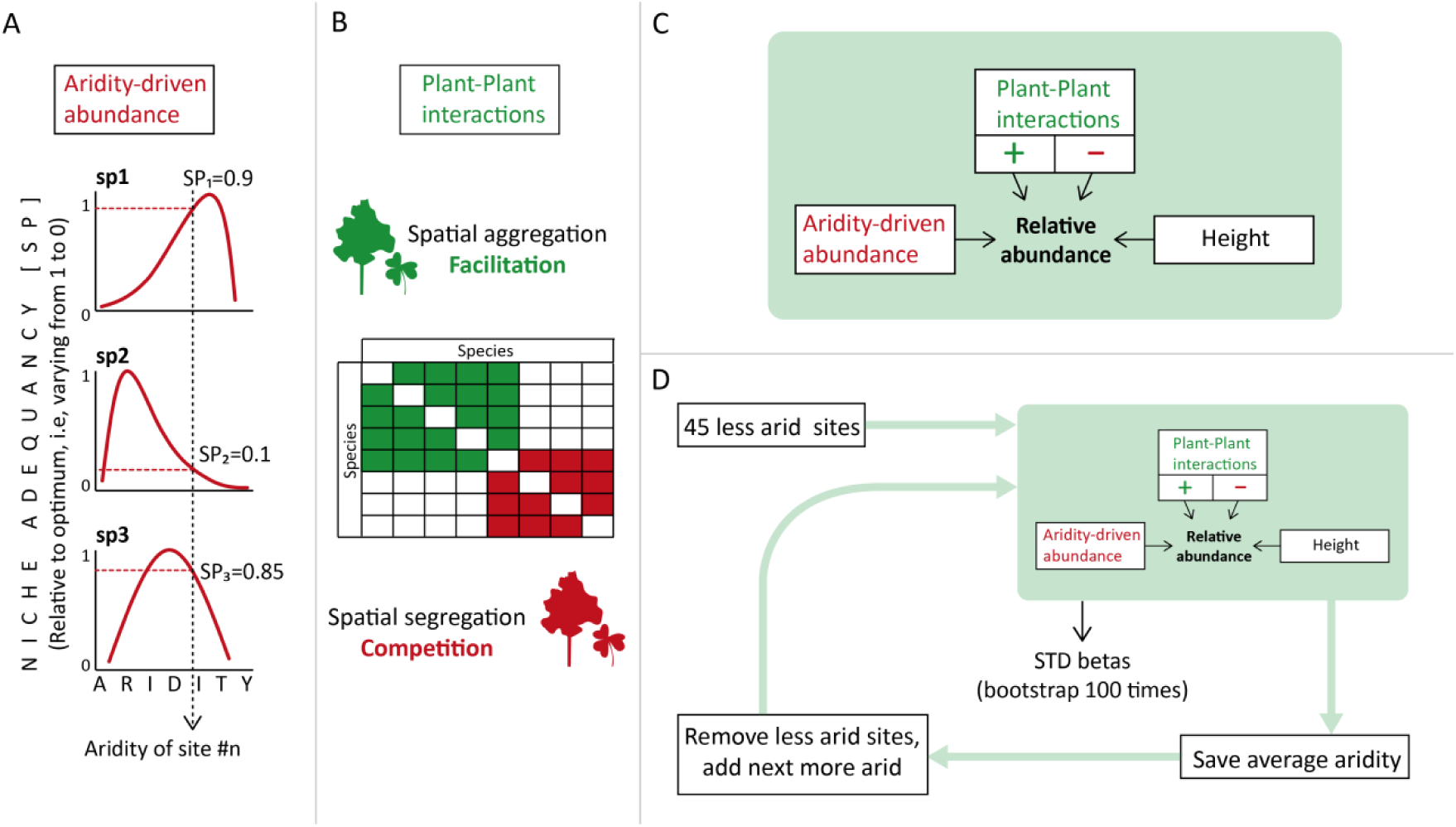
Diagram showing the methodological approach used. First, we obtained, for each species, the Aridity-driven abundance using species distribution modelling with aridity as predictor (a) and spatial co-occurrences measured in the field (as a proxy of plant-plant interactions, b). Then we built the model by additionally controlling for species size and using the relative abundance of species as response variable (c). The effects of these elements on species abundance is a metric on the importance of each assembly driver evaluated (aridity, facilitation and competition) for community assembly. Finally, we used a moving window approach to explore how the importance the different community assembly drivers change along aridity and community specialization gradients (d). SP: Species Performance, STD: Standardized coefficients.

The combination of SDMs and local co-occurrence patterns is gathering increasing attention to assess the relative importance of biotic interactions in species distribution models (e.g., by including co-occurrence matrices on SDMs; Boulangeat et al. 2012; Wisz et al. 2013; Godsoe et al. 2017; or by using joint SDMs with several species; Tikhonov et al. 2017; Staniczenko et al. 2017). Also, information about the niches of species extracted using SDMs has been included in models of local co-occurrences to infer the relative importance of habitat sharing on co-occurrence patterns (Steinbauer et al. 2016). This combination of approaches provides a promising tool to disentangle the relative importance of community assembly factors at community scales. However, such approaches have been rarely integrated into a comprehensive framework and have not been tested against field data covering a wide range of species pools and abiotic conditions. Furthermore, they have not been used to evaluate changes in the importance of biotic interactions along abiotic gradients. The latter would allow the investigation of how interactions between abiotic factors, plant-plant interactions and species-specific responses drive the relative abundance of species at the community scale, something that would provide a solid framework to understand how communities will change the importance of assembly rules with ongoing climate change.

Here we combine SDMs with local co-occurrence data to test the relative importance of abiotic factors, plant-plant interactions and the degree of specialization, as well as their interplay, as drivers of species abundances within communities. We used data gathered from 157 drylands from all continents except Antarctica (Maestre et al. 2012). These ecosystems are constrained by water scarcity (Whitford 2002), and the interplay of aridity with plant-plant interactions (competition and facilitation) largely drives the composition and diversity of their plant communities (Tielbörger & Kadmon 2000; Tirado & Pugnaire 2003; Soliveres & Maestre 2014). Drylands already cover over 45% of terrestrial surface and will expand their global extent by 11–23% by the end of this century (Prăvălie 2016). Hence, understanding how the relative importance of abiotic/biotic assembly drivers changes along aridity gradients is crucial to predict the response of terrestrial ecosystems to ongoing climate change. Specifically, we hypothesized that: i) species become more adapted and specialized to aridity as the latter increases, ii) aridity and plant-plant interactions (both facilitation and competition) interact to drive community assembly, and iii) in communities dominated by species specialized to aridity, facilitation is less important than competition as an assembly driver.

## Material and methods

### Study sites and field sampling

The 157 sites used in this study are a subset of the 230 sites used by Ulrich et al. (2016), and were located in drylands from 19 countries (Fig. S1). The sites surveyed differ widely in their environmental conditions: annual mean temperature, rainfall and elevation ranges are from −1.8 to 27.8 °C, from 67 to 1219 mm, and from 69 to 4668 m.a.s.l., respectively. Our database includes grasslands, shrublands and savannahs, with species richness ranging from 2 to 52 perennial species, and total plant cover ranging from 2 to 82%.

All the sites were surveyed between 2006 and 2013 according to a standardized sampling protocol (see Maestre et al. 2012 for details). In each of these sites, a 30 m x 30 m plot was established and four 30-m long transects were displayed separated by 8 m from each other. We established 20 quadrats (1.5 x 1.5m) along each transect (80 per site) and visually estimated the cover of each perennial plant species, which was used as our surrogate of species abundance in each quadrat. A total of 898 species were identified to the species level. We calculated the aridity level of each site [1 – aridity index (AI), where AI is the precipitation/potential evapotranspiration] from AI obtained from the Global Potential Evapotranspiration database (Zomer et al. 2008), which is based on interpolations provided by WorldClim (Hijmans et al. 2005).

### Evaluating aridity and plant-plant interactions as assembly drivers

We developed a four-step approach to evaluate how abiotic and biotic factors determine species relative abundances in plant communities. First, we extracted the aridity niches of all species surveyed using SDMs, and the local relative abundance of species expected when considering only abiotic conditions. Then, we estimated the main features of species niche (niche optimum, niche breadth and niche skewness), calculated a community-weighted mean of such features, and evaluated their variation along aridity gradients to identify changes in common strategies of species specialization to aridity across environmental gradients. Third, we evaluated the effect of both aridity and plant-plant interactions (as extracted from co-occurrence analyses) on the relative abundance of species within each community. Finally, we evaluated changes in the relative importance of abiotic/biotic assembly drivers along gradients of aridity and of niche specialisation of the species in our communities. These steps are described in detail below.

#### Step i – Assessing aridity niches using species distribution models

SDMs are nonlinear statistical models relating abiotic variables (predictors) with species occurrences (response variable) at regional or global scales. We obtained species occurrences from the Global Biodiversity Information Facility (GBIF, http://www.gbif.org/). For simplicity, we used only aridity as the sole abiotic factor for our SDMs. Aridity is a good proxy for water availability, which is the most influential abiotic factor for plant survival in drylands (Whitford 2002), and is a key determinant of both species interactions and composition in drylands (Callaway 2007; Soliveres & Maestre 2014).

We performed SDMs using MAXENT (Elith et al. 2011) as fully described in Appendix S1. The result of MAXENT is a function relating the suitability of a species with aridity (i.e., the “aridity niche”). Based on the aridity niches, we then estimated the “aridity-driven abundance” (Aab), i.e., the expected local relative abundance of each species based solely on the aridity level of each surveyed site and the other species able to colonize the site (see Fig. 1.a).

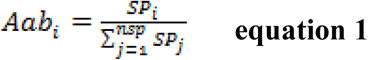

where nsp is the number of species in the community, SP is the species performance calculated as the habitat suitability for the species yielded as an interpolation between the species niche and the aridity observed in the surveyed communities. To ensure comparability of SP, we standardized niches to their maximum (thus it ranged between 0 and 1 for each species). This methodology assumes that, for a particular environmental condition, a species will share the available space with its neighbors by occurring proportionally to its aridity preferences (as measured with the aridity niches). We assumed that relative abundances in the community emerge from sampling the species pool according to the local aridity level, thus, the relative abundance is the density expectation of sampling all species present in the community, each with a probability that depends on its SP.

#### Step ii – Identifying dominant plant strategies based on niche features

Aridity niches hold information about the adaptative strategy of species by showing the following features: i) niche optimum (the aridity level at which a species performs optimally, SP = 1), ii) niche breath (the aridity range that a given species occupies), and iii) niche shape (as measured by the skewness of aridity niches; see examples in Fig. S2). As this information is available for each species, each niche feature can be considered as an attribute of the species related to its response to aridity. These attributes can be used to scale species response to aridity at the community level, i.e. to track how the dominant plant strategy and their diversity within communities change across the global aridity gradient.

First, we calculated the community weighted mean niche optimum obtained as the sum of species niche optimum weighted by their observed relative abundance (adapted from Lavorel & Garnier 2002, hereafter CW-niche optimum) as a measure of the tolerance of communities to aridity. CW-niche optimum was used to evaluate how well the optimum level of aridity of a given species matched with the observed aridity in the surveyed sites. Differences between CW-niche optimum and observed aridity may impact the importance of aridity-driven abundance as a community assembly driver, as it supposes extra stress to species maladapted to local conditions. We used this analysis to understand variations in the importance of aridity-driven abundance (see step iv) across aridity gradients. Additionally, this analysis indicated whether information extracted from SDM matched the one provided by observed patterns, as the local abundance of a species in a given community should exhibit aridity optima around the local aridity conditions of such community (meaning that the community is locally adapted to the aridity).

Second, we calculated the community weighted mean of niche breath (CW-Niche breadth) and shape (CW-Niche skewness) to assess the degree of species specialization to aridity. A smaller niche breath defines species specialized to a particular range of aridity conditions, whereas the shape informs about the preference of such species for more or less arid environments. Hence, communities dominated by species specialized to aridity will be defined by lower CW-Niche breath and negative CW-Niche skewness (i.e., right-skewed, indicating preferences for high aridity level). We observed a strong correlation between niche breadth and skewness (*r* > 0.60): communities dominated by species with a narrow niche breath tend also to be dominated by species with a negative skewness (Fig. 2). Therefore, we used only CW-Niche skewness as a measure of the community specialization towards arid environments (community specialization) in further analyses.

**Figure 2.**
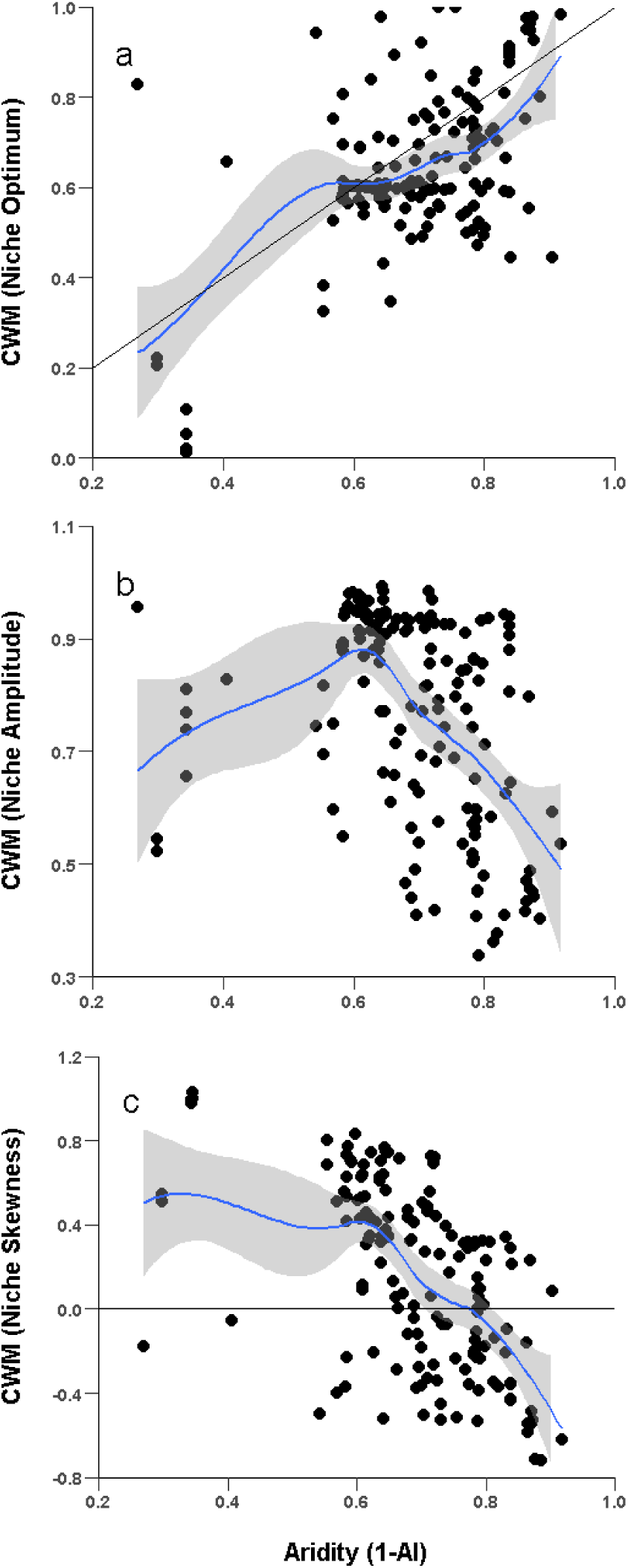
Relationships between aridity and the community weighted mean (CWM) of the niche optimum (a), niche breadth (b) and niche skewness (c) obtained from species distributions models. The blue line and shaded area are the gam-smoothed trends (non parametric regressions) observed ± 95% confidence interval, respectively. The black line in a) represents the 1:1 line; the 0 value in c) indicates a change in the direction of skewness.

#### Step iii – Developping a statistical model to predict species abundance

##### -Plant-Plant interactions: Expected abundance using co-occurrence matrices

For each site, we obtained an estimate of the expected relative abundance of each species according only to plant-plant interactions, measured as spatial co-occurrences. We used aggregation/seggregation as proxies of facilitation/competition, respectively (Tirado & Pugnaire 2003; Cavieres et al. 2006; Valiente-Banuet & Verdú 2008). We are aware that spatial aggregation/segregation can also be driven by other factors such as habitat sharing or seed capture (Morales-Castilla et al. 2015). Interpretation of the results should then consider this limitation; however, co-occurrence has been successfully linked to plant-plant interactions as estimated from manipulative studies (Tirado & Pugnaire 2003), and is the only method available to approximate facilitation and competition at the community level when studying many sites and species (Cavieres et al. 2006; Valiente-Banuet & Verdú 2008).

As a metric of spatial aggregation/segregation, we obtained a normalized score of co-occurrence using PAIRS (Ulrich 2008). PAIRS randomizes the matrices of species occurrences within the quadrats (one per site) and detects deviations from random spatial association patterns in all species pairs while controlling for false positives due to multiple testing (Gotelli & Ulrich 2010). We used an abundance-weighted swap method to randomize species occurrence. This method assumes sampling quadrats with equal probabilities of being colonized and keeps species richness and local abundances constant to account for overall differences in habitat suitability. We obtained co-occurrence in both observed vs. randomized communities for each species pair in each community as:

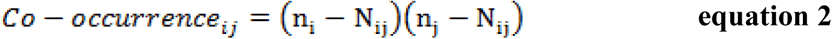

where n is the number of occurrences of target species (n_i_) and its neighbours (n_j_), and is N the number of co-occurrences of both species together. We used the standardized effect sizes obtained from comparing co-occurrences of the null model with that observed in the field as a metric of the strength of the interaction between target species and their neighbours as a function of the deviation from random co-occurrence of species i and j. Thus, standardized effect sizes are comparable between different pairs, but do not take into account how frequent is the interaction within the community. To correct for this, we estimated the relative abundance of a species *i* expected due to competition (i.e., negative co-occurrence, equation 3) and facilitation (i.e., positive cooccurrence, equation 4) with.other species (j, not includying i) as:

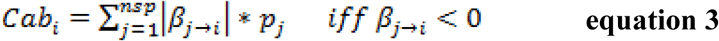

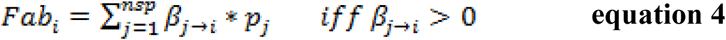

Where, Cab and Fab represent the competition and facilitation-driven abundances in the community; β represents the standardized effect sizes obtained measuring the competitive (if negative) and facilitative (if positive) effect over species *i* of other species in the community (*j*), and pj represents the relative abundance of species *j* in the surveyed community. By doing this we obtained a metric of the effects of plant-plant interactions on the performance of target species for a specific community considering both the strength of interaction with each neighbor (standardized effect sizes) and the frequency of such interactions within the community (relative abundance of the neighbours).

##### -Fitting the statistical model

We used linear mixed models to analyse the relative abundances of each species as a function of: i) aridity-driven abundance, ii) cumulative effects of both competition-driven and facilitation-driven abundances, and iii) the height of the target species (equation 5, Fig. 1.c). Plant height was introduced to control for potential confounding effects between cover (used to estimate relative abundance in the field) and the size of the species being sampled (taller species are more likely to score higher cover values regardless of their abundance). Plant height was obtained from available databases, published literature and local floras (see Appendix A from Soliveres et al. 2014 for a full reference list). Species-specific differences were accounted for by introducing “species identity” as a random factor in the model to avoid the use of the same species in two different communites as two independent cases.

As species relative adaptation to local aridity may influence the importance of facilitation and competition (Choler et al. 2001; Liancourt et al. 2005; Gross et al. 2010; Soliveres et al. 2014), we established an interaction between aridity-driven (derived from the niches and summarizing species suitability to local conditions) and competition- and facilitation-driven abundances. Interactions between aridity and competition and facilitation will be positive if the effect of plant-plant interactions on relative abundance is higher for locally adapted species than for species not adapted to local conditions. It must be noted that the effects of competition are negative, therefore positive contributions from the interaction term decrease the effect of competition on the relative abundance of species adapted to local aridity conditions. Thus, our final model was:

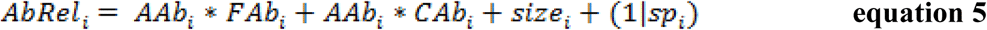

Where *AAb, FAb* and *CAb* represent aridity, facilitation and competition-driven abundances, respectively. Size is the height of species *i*. We obtained the standardized effect sizes of all variables on relative abundance. We assume that the effect size of how suitable the local aridity is for a given species (AAb) and plant-plant interactions on relative abundance represent the relative importance of abiotic factors and plant-plant interactions, respectively, as drivers of the assembly of the communities analyzed.

#### Step iv –Exploring changes in the relative importance of biotic/abiotic assembly drivers across gradients of aridity and community specialization

First, we ordered all sites according to either aridity or community specialization (contingent factors). Then, we took the 45 sites with the lowest values of each contingent factor (as this number of sites allowed sufficient statistical power for our model), and performed the mixed model described in equation 5 (see Fig. 1.d). In the case of community specialization, we did not use the interaction terms described in equation 5. We did so because CW-Niche skewness already summarizes the influence of species adaptation on the importance of plant-plant interactions and, therefore, the information extracted from interaction terms is redundant with that extracted from the gradient.

We bootstrapped the standardized slopes of each predictor to obtain their confidence intervals, which were matched to the average value of the contingent factor studied across the 45 sites. Next, we removed the community with the lowest value of the contingent factor studied from the 45 selected sites, and added the community scoring the next higher value to repeat the same calculations. We repeated this loop as many times as sites remained (112). The coefficients of the standardized predictors included in the linear mixed models provide a comparable measure of the importance of plant-plant interactions and position of each species regarding its aridity niche. We used the 95% confidence interval to assess changes in the importance of biotic/abiotic assembly drivers across the gradients studied.

### Further statistical details

To maintain information representative of the community level in the analyses described in steps ii, iii and iv above, we used all sites for which we gathered enough information (e.g., discarding species with less than 20 occurrences [see appendix S1], or those for which we could not retrieve height values) for the species that summed up at least 60% of the total perennial vegetation. A total of 157 out of the original 236 communities remained for further analyses, leaving a total of 1631 study cases (405 different species in 157 communities with some species repeated throughout communities). The species from these communities represented on average of 91.6 ± 10.3% (mean ± SD) of the total cover in the surveyed sites.

Mixed models in steps ii and iv were performed using the “lme4” R package (Bates et al. 2015) in R (R Development Core Team 2008). We log transformed all variables but aggregation and segregation (which were double square root transformed), and scaled the values after transformation to fulfill the assumptions of the analyses and to obtain standardized coefficients. We extracted the marginal (variance explained by fixed factors) and conditional (variance explained by fixed + random factors) R^2^ values (Nakagawa & Schielzeth 2013) using the “piecewiseSEM” R package (Lefcheck 2015).

In analyses of steps ii, and iv we used generalized additive models (Wood 2006) to depict smoothed trends in the effects of community niche features and assembly drivers across gradients of aridity and community specialization. These models are used to investigate the nonlinear relationships and work well when a large number of replicates is considered (Wood 2006). Data and code used to perform all the analyses are available in figshare (Berdugo et al. 2017a).

## Results

### Common strategies on species adaptive response along aridity gradients

The relationship between the CWM of aridity optima and observed aridity was close to the 1:1 line (slope = 0.8 ± 0.19), but deviated from this line at intermediate aridity levels (about 0.6–0.8; Fig. 2a). Both CW-Niche skewness and CW-Niche breadth decreased within this aridity range, suggesting that species became more specialized to arid conditions by skewing their niches to the right (i.e., showing preference for more arid environments; Fig. 2b and 2c). All these trends were not confounded by the uneven distribution of the number of communities across the aridity gradient (Fig. S3).

### Changes in the relative importance of aridity and plant-plant interactions as assembly drivers

The strongest predictors of the relative abundance of each species were facilitation (measured as positive co-occurrences) and aridity, which exhibited similar effect sizes (Fig. 3). Competition (negative co-occurrences) and the interactions between aridity driven abundance and plant co-occurrences showed negative effects in the overall model. The negative effects of interaction terms suggest that species well adapted to aridity in drylands usually experience less facilitative effects and more competitive effects.

**Figure 3.**
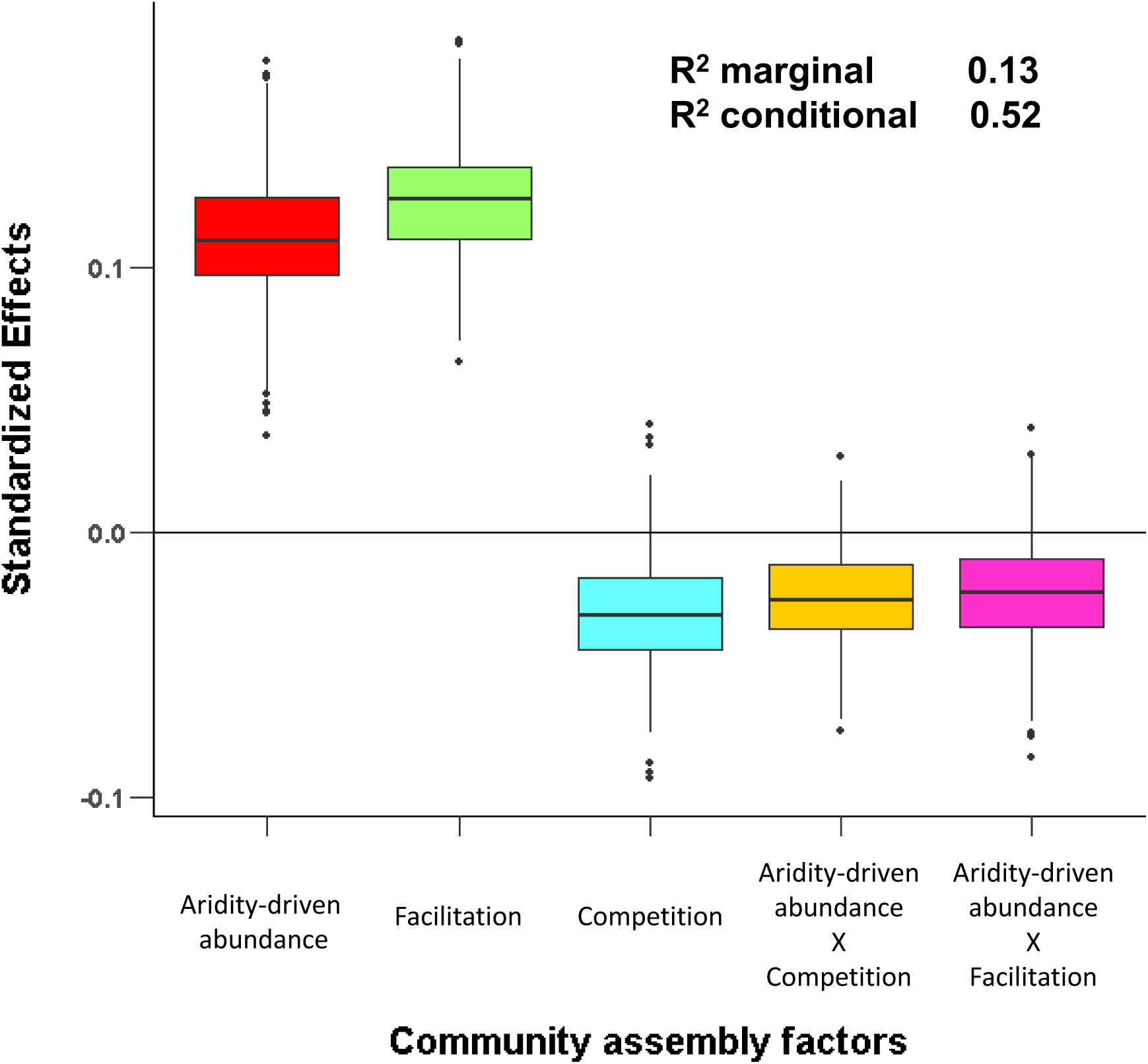
Standardized effect sizes of different drivers of community assembly obtained from the linear mixed model applied to all dryland communities. Median, 50 and 75 quantiles are represented in a box plot for each effect. Marginal (variance explained by fixed factors) and conditional (variance explained by fixed + random factors) R^2^ values are shown.

The importance of aridity as an assembly driver increased up to aridity levels ~ 0.75 (i.e., the limit between arid and semiarid climates), and remained constant beyond that value (Fig. 4a, see also Table S1). The effect of facilitation declined linearly, while that of competition increased (i.e., became more negative), with aridity. However, the effect of competition was only significant under very high aridity levels (0.75–0.80). The interaction term between aridity-driven abundance and competition shifted from negative at wetter sites to positive at dryer sites. These results indicate that, in the less arid sites of our gradient, competition was less important for species more adapted to local aridity than for those less adapted to them. Conversely, at high aridity levels, the effects of competition were stronger for species well adapted to aridity than for those that were far from their aridity optimum. The interaction term between response to aridity and facilitation turned negative (Table S1, Fig. 4b), although only significant in some points of the gradient, with increasing aridity. This result suggests that, under high aridity conditions, facilitation tend to be a more important driver of species abundances for those species maladapted to the observed (high arid) conditions.

**Figure 4.**
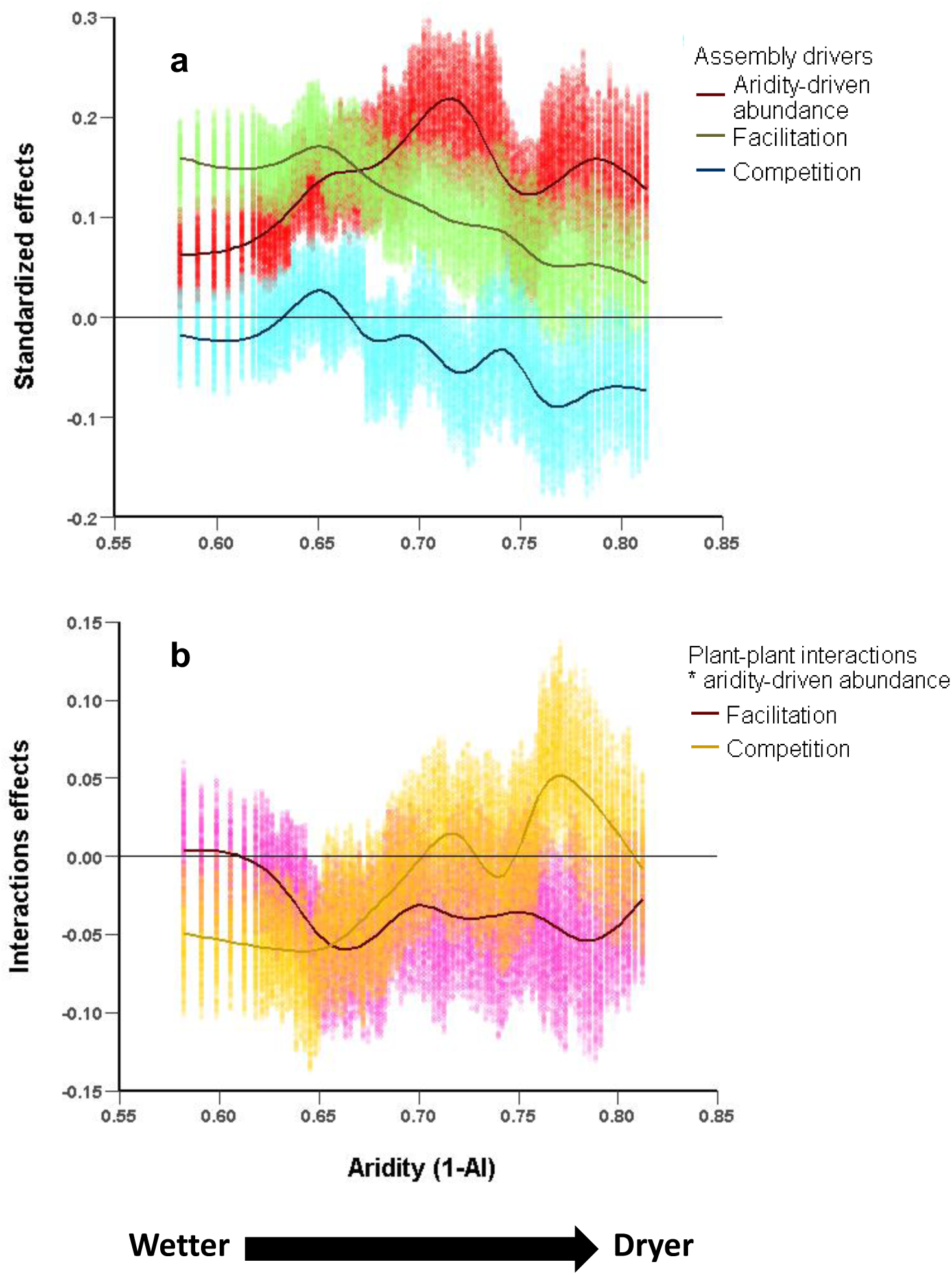
Standardized effects sizes of facilitation, competition and local performance to aridity along an aridity gradient (a), and interactions between local performance to aridity and competition or facilitation (b). This analysis is performed by fitting a generalized mixed model throughout a moving window subsetting our study sites following the gradient of aridity. Bootstrapped coefficients of this regression within the95% confidence intervals are displayed for each step of the moving window. Lines are the gam smoothed trend of variation of the effects.

As communities became more specialized, the importance of facilitation as a community assembly process decreased (Fig. 5). This corresponded with an increase of the importance of competition and aridity-driven abundance. The decline in the importance of facilitation was abrupt and became not significant around values of skewness=0, thus representing Gaussian-like shapes. These results remained consistent when using autoregression analyses instead of generalized additive models (Table S1).

**Figure 5.**
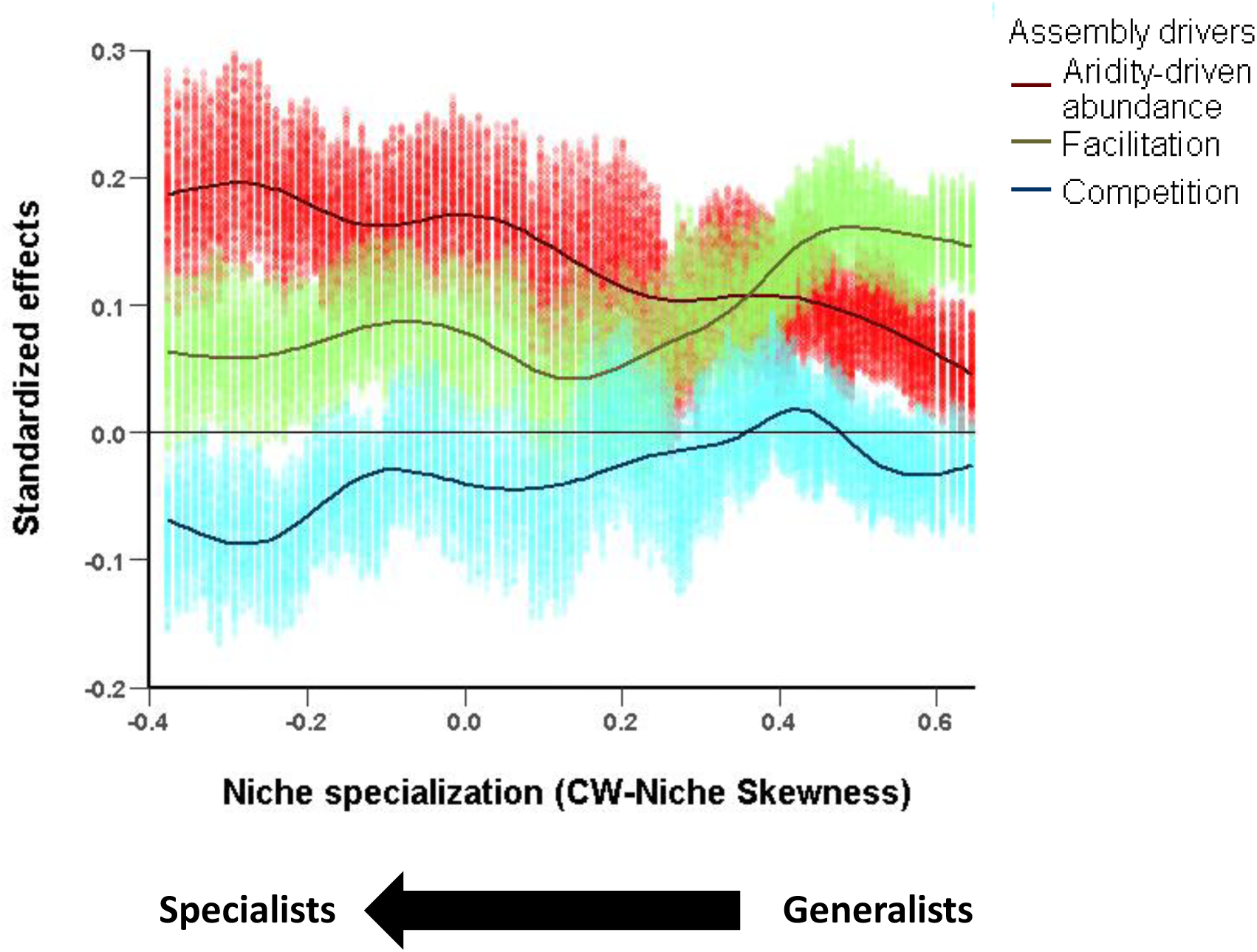
Standardized effects sizes of facilitation, competition and aridity along a gradient of community specialization (measured as community weighted [CW] niche skewness). This analysis is performed by fitting a generalized mixed model throughout a moving window subsetting our study sites following the gradient of community specialization. Bootstrapped coefficients of this regression within the 95% confidence intervals are displayed for each step of the moving window. Lines are the gam smoothed trend of variation of the effects.

## Discussion

By merging approaches operating at contrasting spatial scales, we showed the differential effect of abiotic conditions and plant-plant interactions according to current aridity levels and the degree of habitat specialization of the species pool to aridity. Our study provides fundamental information on how species assemble in global drylands and may help to forecast future community composition in response to climate change. We found that facilitation (measured as positive spatial co-occurrences) and aridity largely explained community assembly in global drylands. The weak effects of competition and the interactions between abiotic and biotic drivers observed were explained by shifts in the importance of abiotic/biotic assembly drivers across gradients of aridity and specialization. We also observed a shift towards communities more specialized with aridity, which substantially reduced the importance of facilitation in the assembly of these communities at the most arid sites studied.

### Facilitation as driver of community assembly in drylands

We found that facilitation and aridity were the two main assembly factors in dryland plant communities. The high importance of facilitation on explaining local plant abundance found here suggests that species niches might be highly influenced by this particular biotic factor in dryland communities. Particularly, the importance of facilitation was maximum in mild drylands (those with aridity values lower than 0.7) and for generalist communities (Fig. 3). Our results also indicate that facilitation importantly affect the abundances of not only rare but also dominant species in these sites, as previously documented (Le Bagousse-Pinguet et al. 2014). This peak in the importance of facilitacion could be driven by two ecological mechanisms: i) species less well-adapted to local conditions are those that benefit the most from facilitation, specially at moderate levels of stress (e.g., Choler et al. 2001; Maestre et al. 2009; Holmgren & Scheffer 2010), or ii) species with extended ranges (and therefore appearing as generalists) in drylands might seem so because positive biotic interactions substantially extend their niches (Wisz et al. 2013; Stewart et al. 2015; Tikhonov et al. 2017). However, with our approach is not possible to distinguish between these two mechanisms. Given that biotic interactions show stronger effects at local scales (Pearson & Dawson 2003), and that the optimum of the niches matches the aridity of our sites (thus suggesting that niches were primarily linked to abiotic conditions, Fig.2), we assume that the former mechanism (facilitation favouring less adapted species) is more likely to occur in the communities studied. However, the high importance of facilitation in our study supposes a good starting point to encourage the study of how facilitation may affect the niches of dryland species at mild environmental conditions.

### Changes in abiotic/biotic controls of community assembly across aridity and specialization gradients

Our SDMs indicated that species niches became narrower and more skewed to dry conditions at aridity levels > 0.7 (Fig. 2). This suggests a high degree of specialization of those species growing under more arid conditions, probably as an adaptive response of communities to increasing environmental harshness (Noy-Meir 1973; Devictor et al. 2010). In contrast to previous predictions that did not account for the degree of specialization of the species pool (Butterfield 2015), our results suggest that communities already experiencing high levels of aridity should not be expected to drastically shift their composition with further aridification. The latter is supported by the asymptotic trend on the importance of aridity-driven abundance in the most arid drylands (Fig. 4a for aridity > 0.75, Fig. 5), and the lower levels of beta-diversity found in these communities (Ulrich et al. 2014). Our results highlight the importance of including the degree of specialization of the species pool to accurately forecast compositional shifts with climate change, as highlighted by Bush et al. (2016).

Concomitantly, specialization of the species pool affected also the relative importance of plant-plant interactions at the community level (Fig. 5, Fig. S5). Indeed, we found a collapse of facilitation as a driver of plant community assembly – as previously forecasted by studies focusing on pairwise interactions (e.g., Tielbörger & Kadmon 2000; Cavieres et al. 2006; Michalet et al. 2006) – to occur at aridity levels of 0.75–0.80, likely because of a higher specialization of the species pool at high aridity levels. The assembly of communities dominated by specialist species is less dependent on facilitation and more on competition, as suggested also by experimental studies (e.g., Liancourt et al. 2005; Gross et al. 2010). Our results, therefore, validate the occurrence of a tight relationship between species specialization and the outcomes of plant-plant interactions within plant communities. As specialists became dominant in high aridity sites (Fig. 2b and c), the community-scale importance of facilitation declined, and that of competition increased, along the aridity gradient evaluated (Fig. 4a). Note that within high aridity sites, facilitation was still more important for species not adapted to local aridity conditions than for those not adapted to them (Fig. 4b). Overall, our results support the notion that facilitation is less important for locally-adapted species, and that plant-plant interactions depend more on species-specific adaptations than on the overall environmental harshness (Liancourt et al. 2005; Soliveres & Maestre 2014).

Using a method based on the abundance patterns of interacting species rather than on species richness, we were able to quantify where the shift on the relative importance of facilitation and competition occurs in drylands, something that happened around aridity levels of 0.75–0.80. Importantly, studies focusing on the relationship between facilitation and species richness that focused on pairings between dominant nurses and their neighbours did not find such facilitation collapse along similar aridity ranges as those studied here (Soliveres & Maestre 2014), although this collapse has been found in studies considering temporal variation in climatic conditions (Tielbörger & Kadmon 2000). This suggests that plant-plant interactions behave differently across environmental gradients depending on whether we focus on particular pairs of species or in all possible pairs within a given community, and also on whether we focus on species richness or changes in relative abundances.

Our results might explain why spatial patterns of dryland vegetation decouple from facilitation under aridity levels ≥ 0.8 (Berdugo et al. 2017b). Under these conditions, facilitation is no longer an important driver of species abundance, which is likely related to the size of plant patches in drylands. Previous studies have failed to link facilitation with ecosystem functioning (Maestre et al. 2010), probably due to the focus on the relationship between facilitation and species richness as the sole mechanism linking facilitation to ecosystem functioning (but see Mitchell et al. 2009). We speculate that focusing on the links between facilitation and species abundance, known to affect spatial patterns that are fundamental drivers of ecosystem functioning in drylands (Maestre et al. 2016), could provide the long hypothesized but largely untested link between facilitation and ecosystem functioning. Interestingly, at approximately the same aridity levels where facilitation declines and communities become more competition-driven, other studies have found important functional changes involving nutrient cycling rates (Wang et al. 2014) and drastic declines in ecosystem functioning (Berdugo et al. 2017b). Since plant-plant interactions are thought to affect ecosystem resilience (Kéfi, Holmgren & Scheffer 2016), our study points that a possible driver of such sensitivity might be a shift from facilitation to competition-driven plant communities.

## Concluding remarks

The SDMs effectively isolated the effect of abiotic conditions, as supported by the good fit of estimated aridity optima of the observed community vs. the local aridity of each community (Fig. 2a). In addition, our results are in accordance to those estimating the relative importance of biotic/abiotic assembly drivers from patterns of height variation (Fig. S4) (Cornwell & Ackerly 2009; Mayfield & Levine 2010). Thus, the combination of approaches at different spatial scales introduced here effectively allowed us to isolate the role of abiotic conditions and plant-plant interactions when analyzing the relative abundance of species within communities. We found shifts from facilitation- to competition-driven communities under aridity levels around 0.75, with potentially important cascading effects on ecosystem functioning that deserve further attention. Furthermore, by explicitly considering the adaptation of species to aridity we showed that facilitation was more important for maladapted species in drylands, and that specialized species pools estabilize the effect of environmental filtering across a large range of aridity levels. Our results emphasize the role of species adaptation to aridity as a modulator of the role of environmental filters and plant-plant interactions as drivers of community assembly. They also suggest that the composition of arid plant communities may be highly resilient to further increases in aridity, and that facilitation is key to preserve species less adapted to high aridity levels. These findings can be used to refine forecasts of plant community composition under climate change in drylands, the largest biome on Earth.

## Acknowledgements

We thank Concha Cano for her help on the illustrations displayed in some figures and all the members of the EPES-BIOCOM network for the collection of field data. This work was funded by the European Research Council under the European Community’s Seventh Framework Programme (FP7/2007–2013)/ERC Grant agreement 242658 (BIOCOM). MB was supported by a FPU fellowship from the Spanish Ministry of Education, Culture and Sports (Ref. AP2010-0759). FTM acknowledges support from a Humboldt Research Award from the Alexander von Humboldt Foundation and from the European Research Council (ERC Grant agreement 647038 [BIODESERT]). The research of SK has received funding from the European Union’s Seventh Framework Programme (FP7/2007–2013) under grant agreement no. 283068 (CASCADE).

## Author’s contribution

F.T.M. designed the sampling design and coordinated field data acquisition. M.B. and S.S. designed the study. Data analyses were done by M.B, assisted by S.S. The paper was written by M.B., and all authors substantially contributed to the subsequent drafts.

## Data accessibility

Dataset used in this study and R codes used to perform analyses are fully available through figshare (Berdugo et al. 2017a).

## Supplementary figures

**Figure S1.**
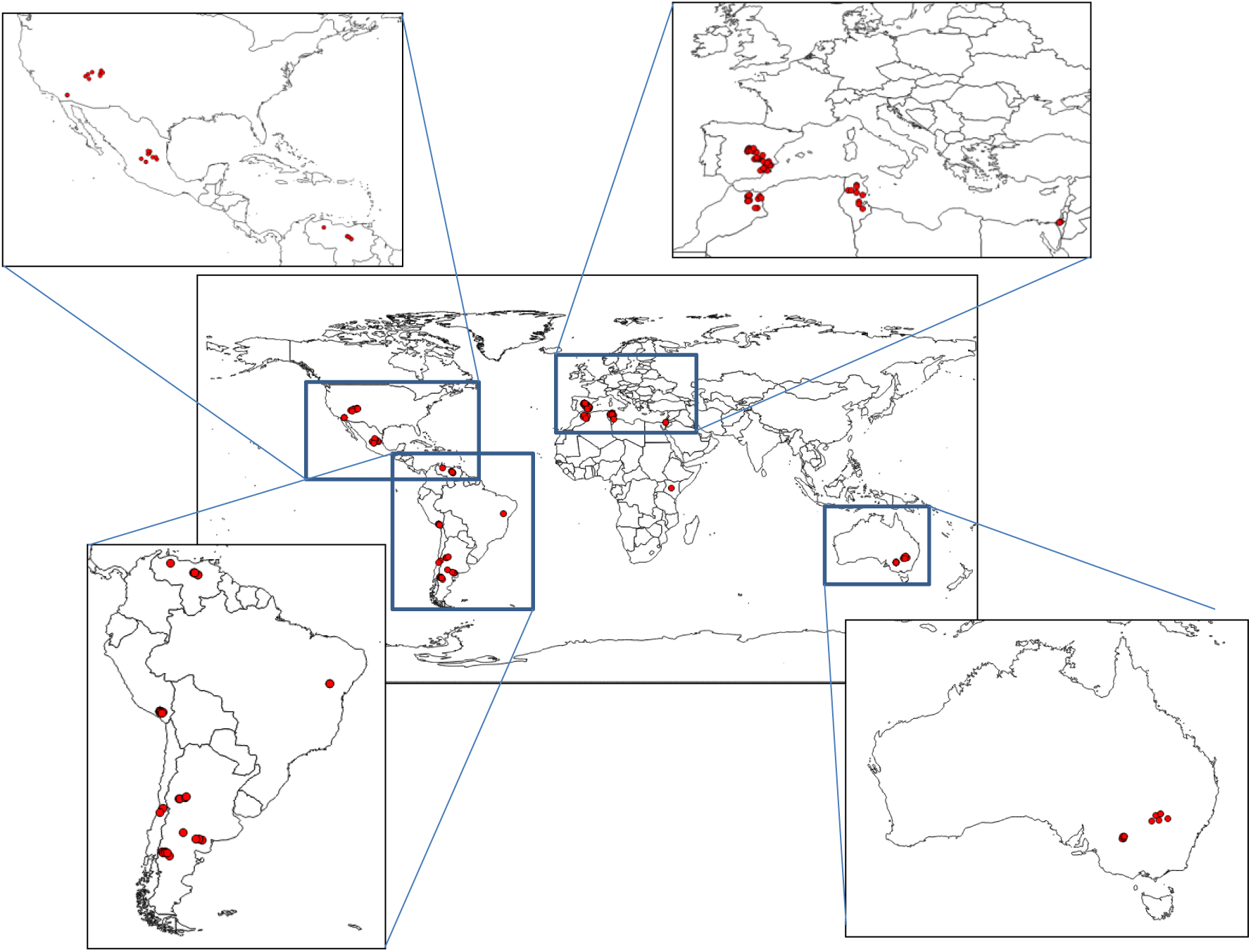
Map showing the geographical position of the 157 sites used in this study (red points).

**Figure S2.**
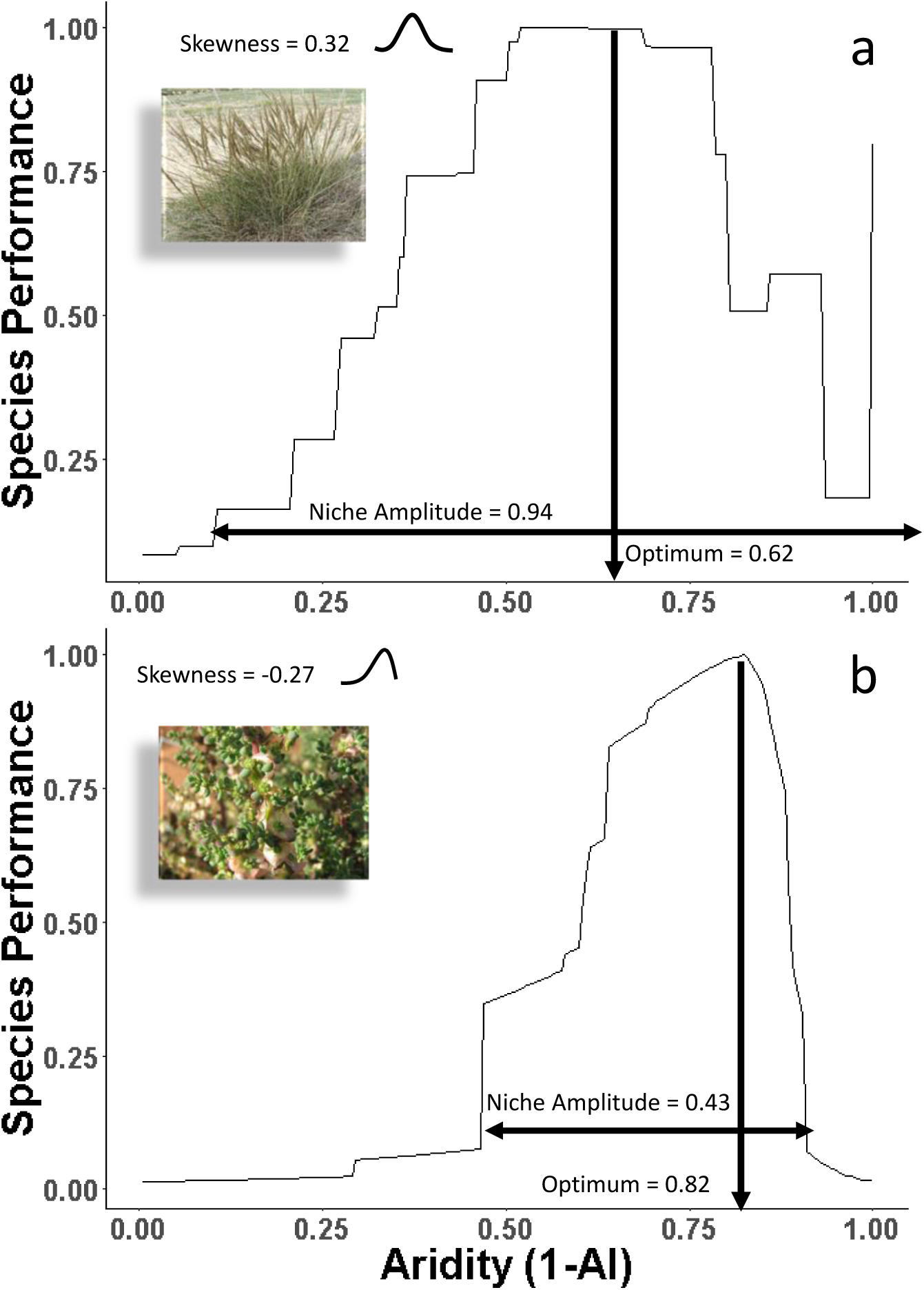
Example of aridity niches for *Stipa tenacissima* (a) and *Maireana brevifolia* (b), including the features measured on them. AI = aridity index. Photographs of each species are shown in the figure. Authors: a) Lumbar~commonswiki; b) BY-SA 3.0, downloaded from Wikipedia under creative commons license.

**Figure S3.**
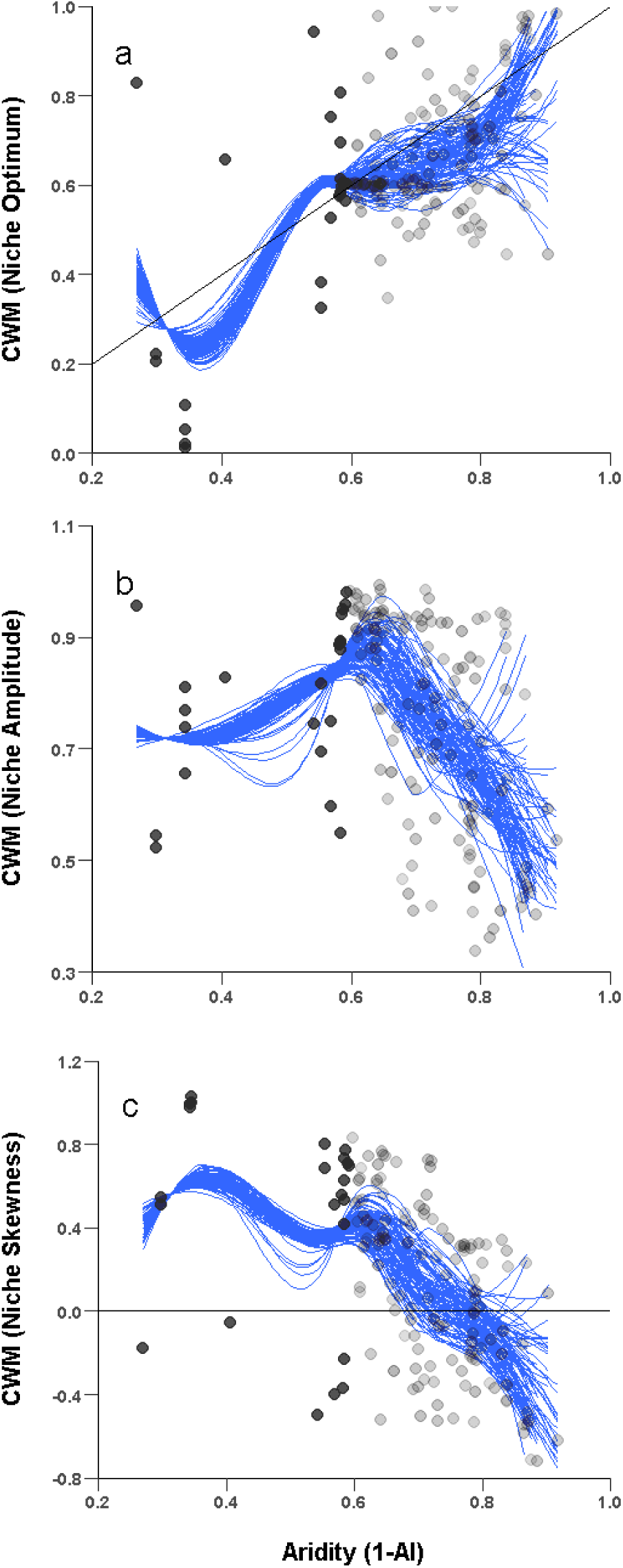
Variation in the relationship between aridity and community weighted niche optimum (a), amplitude (b) and skewness (c) when using the same number of points at both sides of the intermediate aridity level of the study (Aridity = 0.6). We performed this regression 100 times by keeping the sites with aridity lower than 0.6 (N=24) and bootstrapping sites with aridity higher than 0.6 (so that total N = 48 for each regression). Each dot represents a community observed in one of the sites. The transparency of the data points is inverselly proportional to the number of times the point was used. The blue line is the loess smoothed trend observed in each of the 100 bootstrapped samplings. The black line in a) represents the 1:1 line and in c) the 0 value, and indicates a change in the direction of skewness.

**Figure S4.**
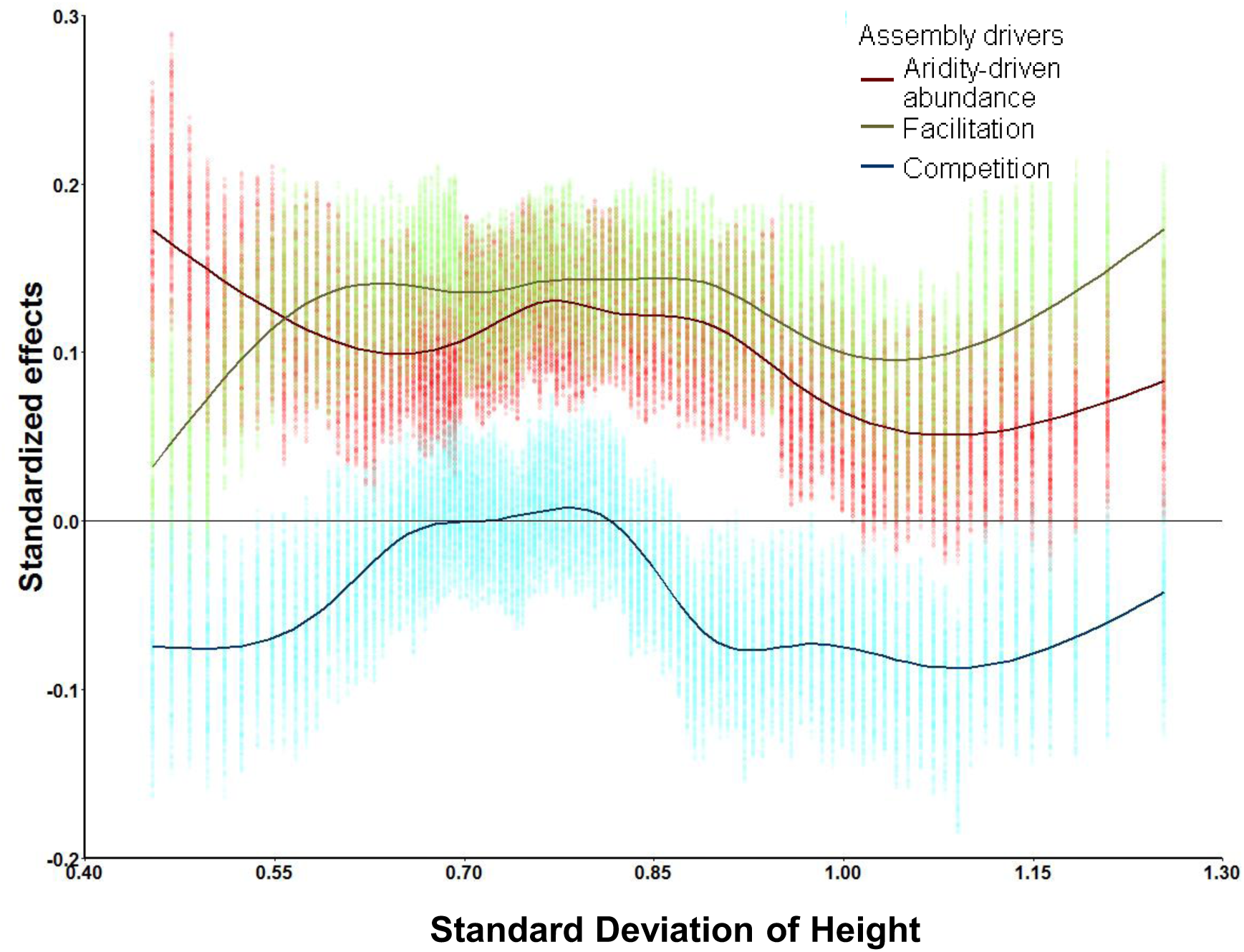
Standardized effects sizes of facilitation, competition and aridity along an aridity gradient. This analysis is performed by fitting a generalized mixed model (see equation 5) throughout a moving window subsetting our study sites following the gradient of height standard deviation within communities. Bootstrapped coefficients of this regression within the 95% confidence intervals are displayed for each step of the moving window. Lines are the gam smoothed trend of variation of the effects.

**Figure S5.**
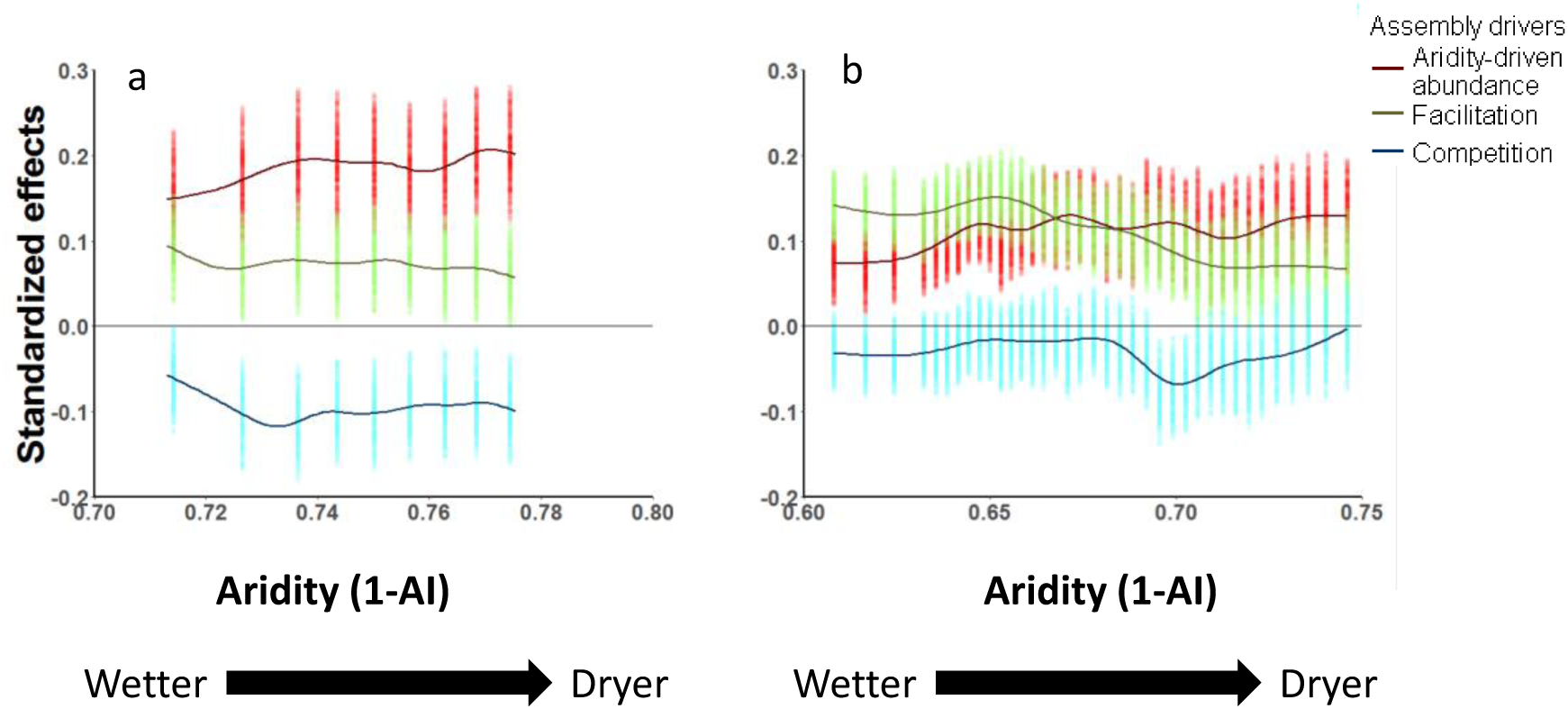
Standardized effects sizes of facilitation, competition and aridity along an aridity gradient for Specialized communities (CWSkewness <0, a), and not specialized communities (CWSkewness >0, b). This analysis is performed by fitting a generalized mixed model throughout a moving window (see figure 1d) in two subsets of our study sites according to CWskewness. Bootstrapped coefficients of mixed models regression within the 95% confidence intervals are displayed for each step of the moving window. Lines are the gam smoothed trend of variation of the effects.

**Table S1.** Results from a lineal and quadratic autorregresive model relating aridity and CW-Niche skewness with the importance of the different drivers of community assembly evaluated. The analysis takes the average of effect sizes per level of the contingent factor (as in Figures 4 and 5) as response variables and uses lagged values of these effects along the gradient at lags = 1 and 2 as covariates of aridity/CW-Niche Skewness to control for autocorrelation. Int. Facil: interaction of Aridity-driven abundance and facilitation. Int. Compet: interaction between Aridity-driven abundance and competition.

|  | Assembly driver | Slope | Quadratic term |
| --- | --- | --- | --- |
| Specialization | Aridity-driven |  |  |
|  | abundance | -0.01<sup>•</sup> | -0.03 (n.s.) |
|  | Facilitation | -0.01 (n.s.) | 0.02 (n.s.) |
|  | Competition | -0.01 (n.s.) | 0.02<sup>•</sup> |
| Aridity | Aridity-driven |  |  |
|  | abundance | -0.01 (n.s.) | -0.50<sup>•</sup> |
|  | Facilitation | -0.16*** | -0.08(n.s.) |
|  | Competition | -0.09* | -0.04 (n.s.) |
|  | Int. Facil | 0.00 (n.s.) | 0.15 (n.s.) |
|  | Int. Compet. | 0.82<sup>•</sup> | -0.55<sup>•</sup> |

Note: \*\*\* significant at P<0.001, \*\* significant at P<0.01, \* significant at P<0.05, <sup>•</sup> marginally significant, (n.s) not significant

